# Zinc-based Ultrasensitive Microscopic Barrier Assay (ZnUMBA): a live-imaging method for detecting epithelial barrier breaches with spatiotemporal precision

**DOI:** 10.1101/2022.09.27.509705

**Authors:** Tomohito Higashi, Rachel E. Stephenson, Cornelia Schwayer, Karla Huljev, Carl-Philipp Heisenberg, Hideki Chiba, Ann L. Miller

**Affiliations:** Department of Basic Pathology, Fukushima Medical University, Fukushima 960-1295, Japan; Department of Molecular, Cellular, and Developmental Biology, University of Michigan, Ann Arbor, MI 48109, USA; Institute of Science and Technology Austria, Klosterneuburg, Austria

## Abstract

Epithelial barrier function is commonly analyzed using transepithelial electrical resistance (TER), which measures the ion flux across epithelia, or by adding traceable macromolecules to one side of the epithelium and monitoring their passage to the other side. While these methods effectively measure changes to global barrier function, they are not sensitive enough to detect local or transient disruptions in the barrier, and they do not reveal the location of barrier breaches within the context of cell or tissue morphology. Therefore, we developed a method that we named Zinc-based Ultrasensitive Microscopic Barrier Assay (ZnUMBA), which overcomes these limitations, allowing for detection of local tight junction (TJ) leaks with high spatial and temporal resolution (Stephenson et al., 2019; Varadarajan et al., 2021). Here, we present expanded applications for ZnUMBA. First, we show that ZnUMBA can be used in *Xenopus* embryos to measure the dynamics of barrier restoration and actin dynamics following laser injury of the junction. We also demonstrate that ZnUMBA can be effectively utilized in developing zebrafish embryos as well as cultured monolayers of Madin-Darby Canine Kidney II (MDCK II) epithelial cells. ZnUMBA is a powerful and flexible method that, with some optimization, can be applied to multiple systems to measure dynamic changes in barrier function with spatiotemporal precision.

## Introduction

Epithelial tissues separate internal compartments within an organism and serve as barriers to the external environment. Formed by sheets of polarized cells connected to one another by cell-cell junctions, epithelia regulate selective transport of ions and solutes across the tissue – either through the cells themselves (transcellular transport) or through the space between cells (paracellular transport). TJs, which are comprised primarily of claudin family and occludin transmembrane proteins along with cytoplasmic linker proteins that connect the TJ to the actomyosin cytoskeleton, are responsible for regulating paracellular transport (Quiros and Nusrat, 2014; Turner et al., 2014; Van Itallie and Anderson, 2014; Varadarajan et al., 2019; Zihni et al., 2016). Along the length of bicellular TJs, claudin-based TJ strands interact between neighboring cells, bringing the cell membranes into close apposition (Farquhar and Palade, 1963; Furuse et al., 1998). The extracellular loops of claudins can form size- and charge-selective pores, where some claudins are predominantly pore forming toward cations or anions, while others are predominantly barrier forming (Gunzel and Yu, 2013). At tricellular TJs (tTJs), where three cells come together, specialized proteins including tricellulin and angulins help to seal the paracellular space (Higashi and Chiba, 2020; Higashi and Miller, 2017).

The size- and ion-selectivity of TJs differs in different epithelial tissues, and can be modulated by both physiological and pathological stimuli. For example, changes in TJ protein expression or post-translational modifications (Bolinger et al., 2016; Gunzel and Yu, 2013; Raleigh et al., 2011), TJ strand number or degree of branching (Claude and Goodenough, 1973; Saito et al., 2021), inflammatory stimuli (Ivanov et al., 2010; Luissint et al., 2016), or tissue mechanics (Stephenson et al., 2019; Varadarajan et al., 2019; Varadarajan et al., 2021) can all modulate barrier function. Therefore, researchers often seek functional readouts to measure the permeability characteristics of epithelia. Traditionally, this is achieved with a combination of two key assays: TER and tracer permeability assays (Matter and Balda, 2003) (**Fig. 1A-B**). TER measures ion permeability by generating an electrical current across an epithelial monolayer grown on a permeable support and measuring the resistance (Srinivasan et al., 2015). TER is useful for measuring the magnitude of ion flux across an epithelial monolayer through the pores generated by claudins (Shen et al., 2011; Suzuki et al., 2014). Thus, epithelia with a high proportion of barrier-forming claudins or perturbations that promote improved barrier function will have higher TER (i.e., reduced ion permeability). In contrast, tracer permeability assays measure size-selective permeability characteristics of an epithelial monolayer by adding fluorescent tracer macromolecules of different sizes (e.g., fluorescein dye (332 Da), noncharged polyethylene glycol (PEG) oligomers (e.g., PEG3: radius = 2.8 Å, PEG25: radius = 7 Å), or fluorescent dextrans (e.g., 4kDa, 25kDa, 75kDa, 250 kDa) to the apical side of an epithelial monolayer grown on a permeable support and measuring how much of the tracer passes to the basal side (Matter and Balda, 2003; Van Itallie et al., 2008). In this way, increased Papp (apparent permeability), measured by increased passage of tracer to the basal side, indicates increased macromolecular permeability.

**Figure 1.**
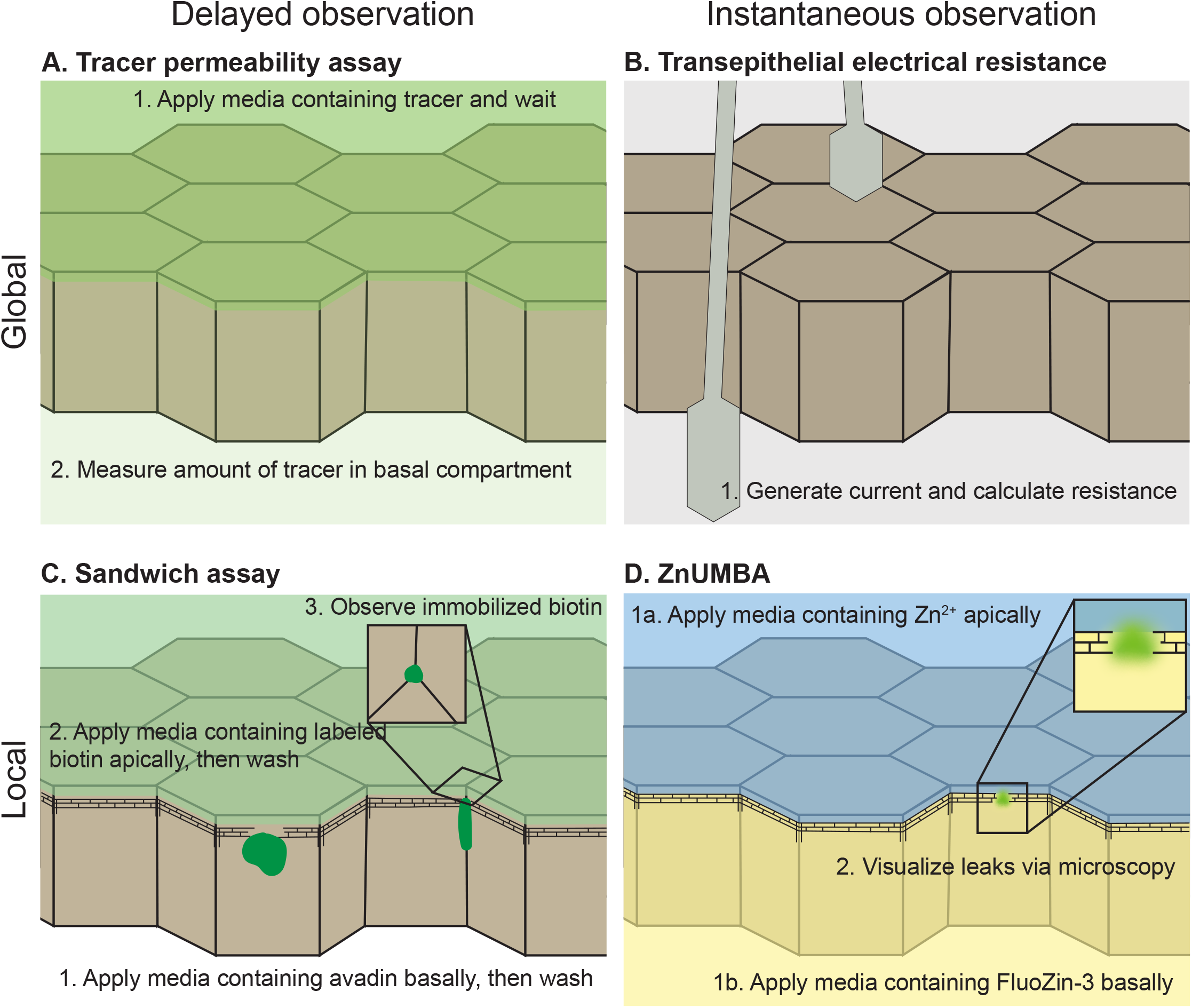
Established methods for assessing epithelial barrier function. Methods for measuring epithelial barrier function can be categorized as measuring global barrier function (top row) or local barrier function (bottom row). Results may be detected after a set waiting time, typically ranging from minutes to hours (left), or in real time (right). **A.** Tracer permeability assays track the passage of macromolecules from the apical compartment to the basal compartment after a set period of time. Permeability measurements are averaged across the whole tissue. **B.** Transepithelial electrical resistance measures the electrical resistance of a monolayer. Measurements can be recorded instantaneously. **C.** Sandwich assays immobilize tracers once they cross the barrier, allowing sites of barrier breaches to be detected at the experiment end point. **D.** ZnUMBA employs the fluorogenic dye FluoZin-3, which increases in fluorescence at sites of barrier breaches, allowing real time monitoring of barrier breaches and their repair.

Together, TER and tracer permeability assays have been the standard approaches to measure the permeability characteristics of cultured epithelial monolayers. However, these methods have a number of limitations: 1) they are global measures that average barrier function across many cells and are not sensitive enough to detect subcellular changes in barrier function within a monolayer, 2) they do not provide spatial information about the location of sites of increased permeability, 3) although they can be carried out over a timecourse, they do not provide information about rapid, dynamic changes in the barrier, and 4) they are not compatible for use with live developing embryos such as *Xenopus*, zebrafish, or mouse embryos.

Several recent methods have sought to overcome these limitations, particularly with respect to identifying the location of barrier breaches within an epithelial tissue. For example, “sandwich assays” allow visualization of local barrier function at fixed time points (Dubrovskyi et al., 2013; Ghim et al., 2017; Richter et al., 2016) (**Fig. 1C**). These assays rely on the high-affinity interaction of avidin and biotin and improve upon previous methods using Sulfo-N-Hydroxysuccinimide-Biotin (Sulfo-NHS-biotin), which binds to primary amines that can be detected by avidin probes (Merzdorf et al., 1998; Schumann et al., 2012; Tamura et al., 2008). Sandwich assays employ, for example, fluorescently-labeled avidin applied to the basal side and fluorescent biotin applied to the apical side – sandwiching the epithelial monolayer. If there is a leak in the barrier, the fluorescent biotin crosses the TJ and is captured by the avidin, and this local fluorescence intensity can be visualized via microscopy at the experimental endpoint. Sequential round(s) of washing the monolayer and adding a different fluorescent biotin can be carried out to visualize snapshots in time, thus revealing changes in the barrier over time. Variations on the “sandwich assay” approach have shown that barrier function to macromolecules is not uniform across the tissue; barrier breaches open and close over time; and changes in barrier function can occur in response to mechanical cues (Dubrovskyi et al., 2013; Ghim et al., 2017; Richter et al., 2016). However, while able to capture local breaches in the barrier, these assays still only reveal snapshots in time. The low temporal resolution of these techniques makes it impossible to investigate the events immediately preceding and following leaks in barrier function. Additionally, multiple fluorescent channels must be used for sequential labeling, reducing the channels available to monitor other fluorescent probes of interest. Thus, sandwich assays provide an advantage over the standard TER and tracer permeability assays in that they can reveal the location of barrier breaches within an epithelial tissue. However, the lack of an assay that is sensitive enough to detect when and where breaches in barrier function happen, how long they last, and whether they repeat at the same sites remains a limitation for researchers investigating dynamics of barrier function.

We developed a method that meets this need: Zinc-based Ultrasensitive Microscopic Barrier Assay (ZnUMBA) (**Fig. 1D**). This method relies on a commercially-available, small, cell-impermeable dye called FluoZin-3 (FZ3, 847 Da, Thermo Fisher Scientific, Cat. F24194). FZ3 is fluorogenic, increasing in fluorescence intensity more than 50-fold when bound to zinc (65 Da); as such, there is little background signal until the dye is bound to its target. Media containing ZnCl_2_ is applied to the apical side of the epithelium, and FZ3 is introduced to the basal side of the epithelium. When localized TJ breaches occur, zinc and FZ3 interact, resulting in bright, localized increases in FZ3 – specifically at the site of TJ damage. Once the leak is repaired and barrier function is reinstated, FZ3 levels return to background levels. Therefore, ZnUMBA can detect local changes in barrier function and examine the dynamics of barrier function with high temporal and spatial resolution using conventional confocal microscopy.

Using ZnUMBA in *Xenopus laevis* embryos allowed us to visualize naturally-occurring, transient leaks in barrier function and correlate these with local loss of TJ proteins, local Rho activation, and actomyosin-mediated repair of barrier function (Stephenson et al., 2019). Furthermore, we recently used ZnUMBA to demonstrate that mechanosensitive channel-mediated calcium influx is required to maintain barrier integrity by repairing local leaks that occur when junctions elongate (Varadarajan et al., 2021). Moreover, Chan et al. (2019) modified ZnUMBA for use in mouse embryos, demonstrating that reduced cortical tension leads to disrupted barrier function of the mouse blastocyst cavity (Chan et al., 2019) and highlighting the versatility of this method for use in other model organisms. Here, we further demonstrate ZnUMBA’s usefulness in *Xenopus* embryos (**Fig. 2A**) and expand ZnUMBA’s application to other systems including zebrafish embryos (**Fig. 2B**) and cultured epithelial monolayers (**Fig. 2C**).

**Figure 2.**
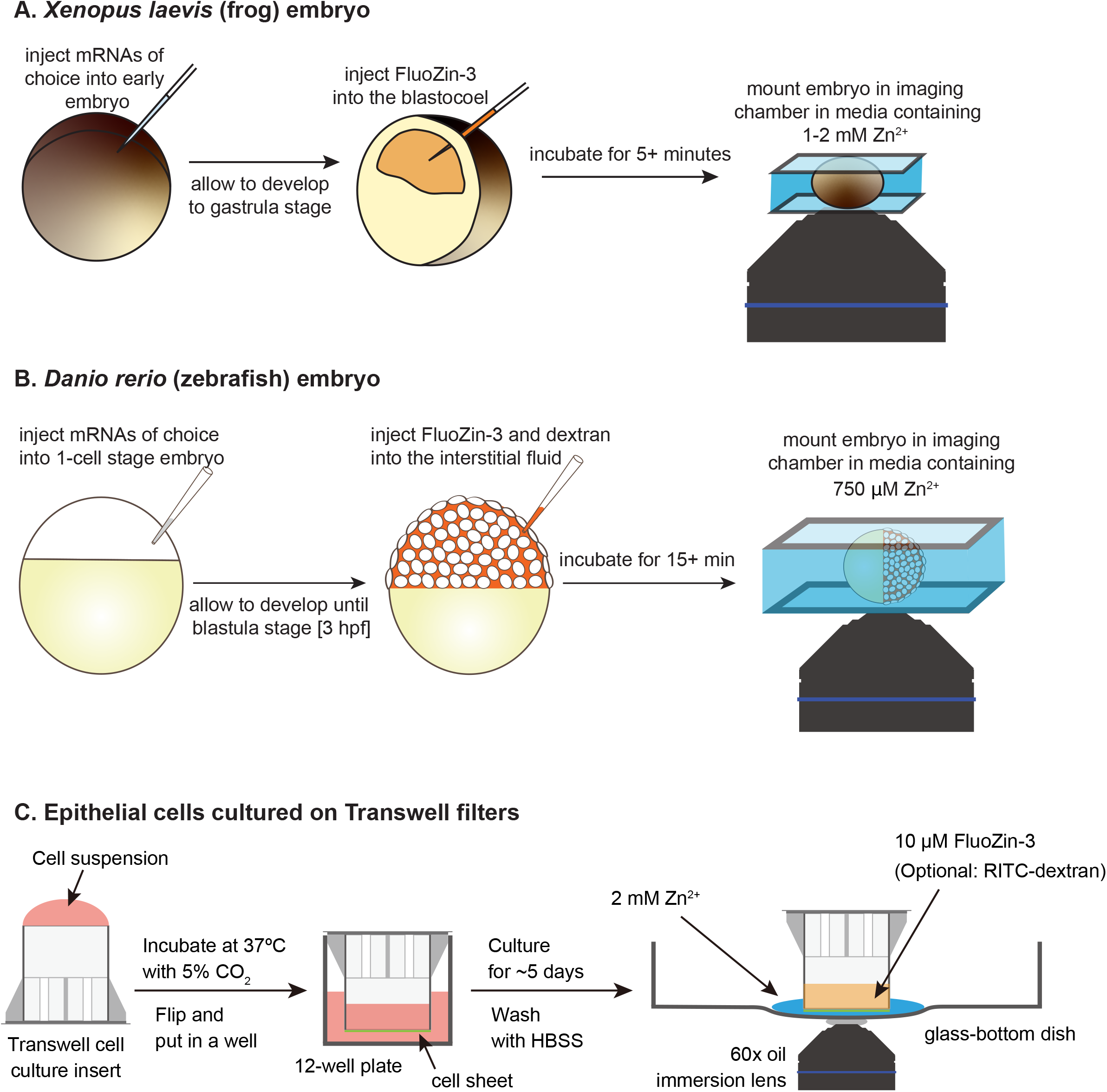
Modification of ZnUMBA for different model systems. **A.** *Xenopus laevis* embryos are injected with mRNAs of interest at early embryo stages (1-4 cell stage). After developing to gastrula stage, FZ3 is injected into the blastocoel, and the embryos are incubated for a minimum of 5 minutes to allow the injection site injury to heal. Finally, just prior to imaging, embryos are mounted in zinc-containing media and imaged via confocal microscopy. **B**. Zebrafish embryos are injected with mRNAs of interest at thw 1-cell stage. Embryos are dechorionated and after developing to 3 hpf (1k-cell stage), embryos are injected with FZ3 and dextran into the interstitial fluid. Embryos are incubated for around 15 minutes to allow injection site injury to heal. Finally, embryos are mounted in low-melting point agarose, and zinc-containing media is added before the start of imaging. **C**. Experimental setup for ZnUMBA using MDCK II cultured epithelial cells. The transwell filter cup is placed upside-down on a clean surface, and 1 x 10^5^ cells resuspended in 300 μL of DMEM are seeded onto the bottom surface of the filter. The filter is incubated at 37°C in a moist CO_2_ incubator for 10-14 hr. After cells are attached to the surface, the filter cup is inverted and placed into a well of a 12-well plate. The cells are cultured for about 5 days until the TER increases. For ZnUMBA, 500 μL of HBSS containing 2 mM ZnCl_2_ is placed on the glass-bottom dish, and the transwell filter cup with the cell sheet attached is placed onto the zinc-containing media. HBSS containing 10 μM FluoZin-3 and 1 μM CaCl_2_-EDTA is added into the filter cup (upper compartment), and the fluorescence is observed by an inverted fluorescence microscope. For visualization of the basal compartment, RITC-dextran could be included in the ZnCl_2_ solution.

## Results

While we designed ZnUMBA primarily to detect local changes in barrier function, we have also demonstrated that ZnUMBA can be used to measure changes in global barrier function. For example, when *Xenopus* embryos were treated with EGTA, junctions become leaky due to chelation of calcium from junctional complexes, which disassembles cadherin-based adhesions, upon which TJs depend (Pitelka et al., 1983; Rothen-Rutishauser et al., 2002; Wang et al., 2016). Similar global disruptions of barrier function were observed when *Xenopus* embryos were treated with SMIFH2, a pan-formin inhibitor (Stephenson et al., 2019), or GsMTx4, a mechanosensitive channel blocker (Varadarajan et al., 2021), suggesting that these treatments result in general TJ structural defects that make TJs universally leakier.

We previously demonstrated that ZnUMBA detects naturally-occurring leaks within elongating junctions, as well as those induced by laser injury of the junction (Stephenson et al., 2019). In both cases, leaks are followed by robust accumulation of active RhoA and actomyosin, which are required for sustained barrier reinforcement (Stephenson et al., 2019). Interestingly, inhibiting the actomyosin response did not result in leaks of longer duration, but in leaks that repeated at the same location over time (Stephenson et al., 2019). In order to more closely investigate this phenomenon, we injected FZ3 into blastocoel of gastrula-stage *Xenopus* embryos as previously described (**Fig. 2A**) (Stephenson et al., 2019), but we refined our laser injury technique and used ZnUMBA to analyze the dynamics of barrier restoration and actin dynamics with higher spatiotemporal resolution (**Fig. 3**).

**Figure 3.**
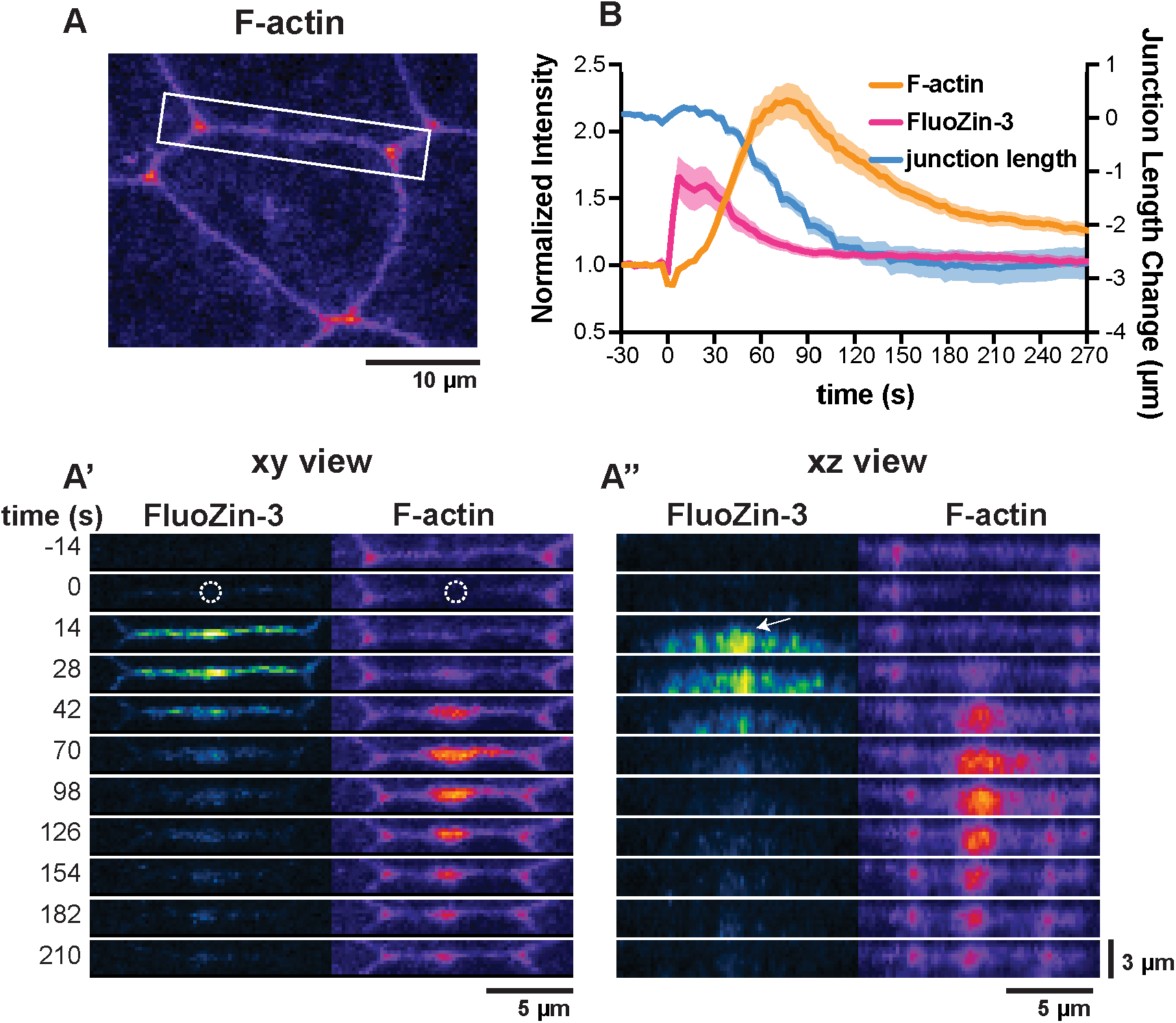
Laser injury induces short-lived barrier breaches in *Xenopus laevis* embryos. **A-A”**. Junction injury was performed by exposing the area indicated by the white dotted circle to intense 405 nm laser light. The junction that was injured is indicated by white box in **A**. Following laser injury (**A’**, time 0), FZ3 (Green Fire Blue Lookup Table (LUT)) increases sharply at the site of injury and less intensely along the length of the junction. F-actin (Lifeact-mRFP, Fire LUT) accumulates at the site of the injury, and the barrier breach is repaired. **A’’** shows side views (x-z) of the images shown in **A**’. White arrow points out that FZ3 signal at the site of the injury is more apical than the signal along the length of the junction **B**. Quantification of laser injury experiments. The mean pixel intensity of a 1 μm wide line drawn over the injured junction from vertex to vertex was normalized to a reference junction from the same movie. Graph shows mean normalized intensity (left axis) or mean junction length change (right axis) ± standard error of the mean (n = 25 junctions, 9 embryos, 3 experiments).

Following laser injury, we observed a sharp spike in FZ3 at the site of the injury as well as less intense signal increase along the length of the junction (**Fig. 3A-A’**). Examining z-views revealed that the FZ3 at the site of the injury is more apical than the signal along the length of the junction (**Fig. 3A”**), leading us to speculate that the lateral signal represents diffusion of FZ3 from the injury site, rather than disruption of TJ function along the length of the injured junction. Furthermore, we observed that FZ3 intensity peaks 10 seconds following laser injury and begins to diminish 30 seconds post-injury, whereas F-actin has only reached ~15% of its maximum intensity by 30 seconds and doesn’t peak until approximately 75 seconds after injury (**Fig. 3B**). Similarly, junction contraction is initiated approximately 30 seconds after injury (**Fig. 3B**). Thus, while actomyosin-mediated contraction appears to be important for sustained barrier restoration, these data indicate that a still unknown mechanism may be able to temporarily seal the barrier prior to full contraction of the junction. While the Rho-mediated response is clearly important for sustained repair, this experiment further highlights that examining barrier leaks with high spatiotemporal resolution can provide new insights into the dynamic nature of epithelial barriers.

ZnUMBA requires that embryos be incubated in micromolar to millimolar concentrations of zinc, which has the potential to interfere with normal cellular function and/or development. In order to test how exposure to zinc impacts overall *Xenopus laevis* embryo viability, we exposed gastrula-staged *Xenopus* embryos to varying concentrations of zinc and assessed gross defects at 1,3, and 24 hours post exposure (**Fig. S1**). After 1 and 3 hours of zinc exposure, abnormalities were rare and appeared at rates similar to controls. However, 24 hours of exposure to zinc did substantially impact development and embryo viability, particularly at the high end of the concentration range tested. Thus, the zinc concentration and duration of exposure should be empirically determined for different model systems.

In order to test the applicability of the ZnUMBA assay in other model systems, we turned to zebrafish embryos. First, early zebrafish embryos were exposed to Danieau’s medium supplemented with different ZnCl_2_ concentrations (**Fig. S2**). Assessing viability at different stages during early development allowed us to identify an optimal condition of 750 μM ZnCl_2_ in which viability largely resembled the control condition (Danieau’s medium only) (**Fig. S2**, red rectangle). Next, we tested the sensitivity and functionality of the ZnUMBA assay in early zebrafish development by artificially inducing barrier breaches through pharmacological or genetic means. We first tested the effect of EGTA addition. Zebrafish embryos were injected with FZ3 into the interstitial fluid, facing the basal side of the outermost epidermal layer (**Fig. 2B**), and at 4 hours post fertilization (hpf), 10-20 mM EGTA (calcium-free medium) was added. The majority of embryos started to become leaky within 90 minutes after EGTA addition, as indicated by the increase in FZ3 intensity normalized to fluorescent Dextran intensity (**Fig. 4A-A’**, FZ3/Dex Ratio). ZnUMBA, therefore, allows for monitoring junctional leakage in zebrafish embryos at high spatiotemporal resolution (see propagation of barrier breaches in **Movie 1**, EGTA lower panel). Next, we sought to induce barrier breaches though a genetic approach, utilizing the *poky* mutant, in which embryos fail to form a functional epidermal barrier due to continued cell cycle and lack of differentiation (Fukazawa et al., 2010). Indeed, when we performed ZnUMBA in *poky* mutant embryos, the FZ3/Dextran Ratio increased withing 80 minutes after starting the assay (Fig. 4 B-B’), indicating that the *poky* mutants did not have a functional epidermal barrier, consistent with previous reports (Fukazawa et al., 2010), and the embryos gradually started lysing around 5-6 hpf. Taken together, these results demonstrate that ZnUMBA can successfully detect barrier breaches in zebrafish embryos due to acute (EGTA) as well as gradually-acquired (*poky* mutant) junctional leakages.

**Figure 4.**
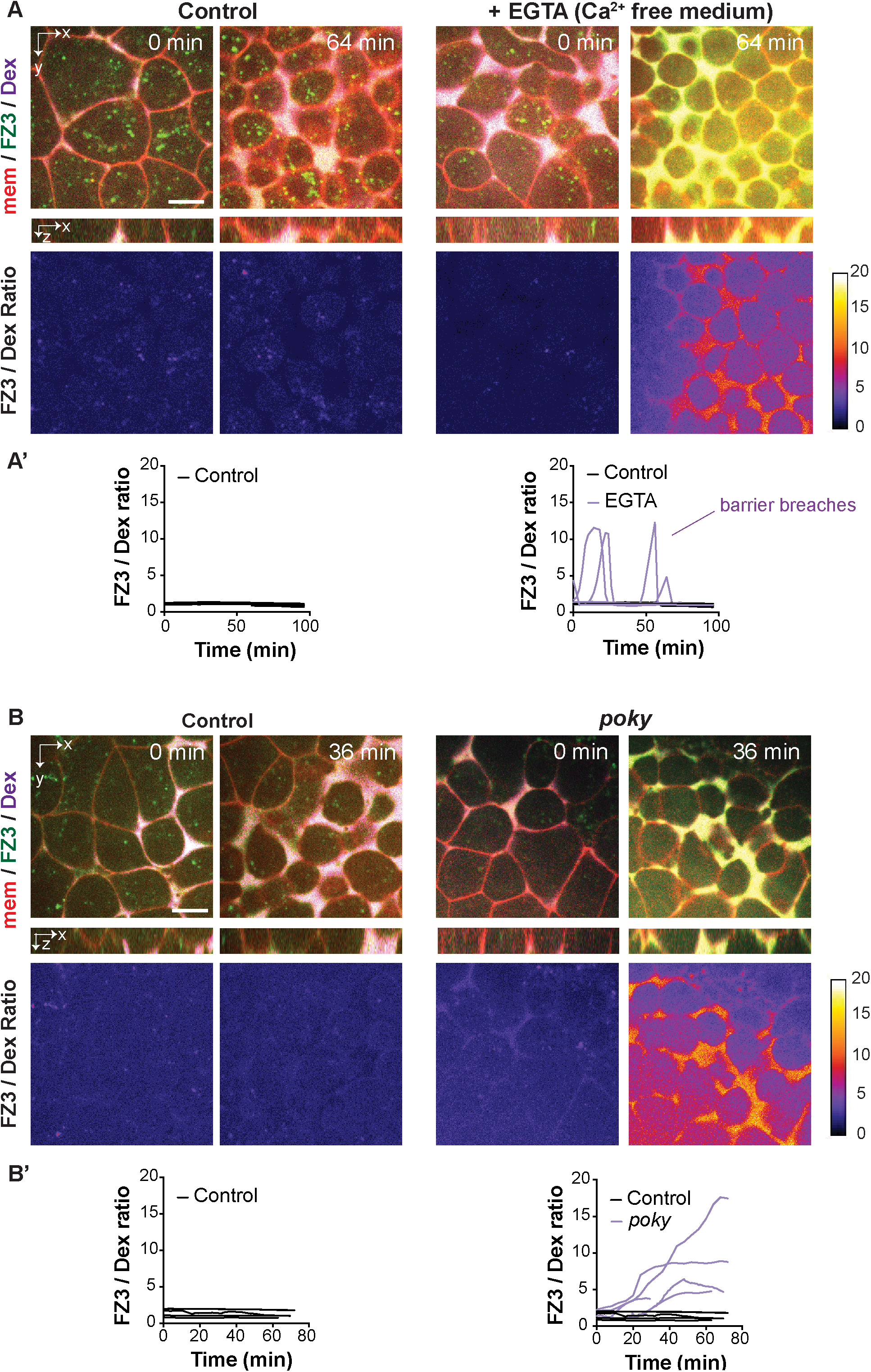
ZnUMBA detects barrier breaches in zebrafish embryos upon pharmacological and genetic perturbations. **A-B.** Consecutive images of an imaging plane at the interface of EVL and deep cells are depicted, with cell membranes in red, FluoZin-3 in green. and Dextran-647 10,000 MW in magenta. Control and EGTA-treated embryos are shown in **A**, while Control and *poky* mutant embryos are shown in **B**. Second row shows side views (x-z) with most interstitial fluid accumulation below the EVL cells. Third row displays the ratio of mean intensities of FluoZin-3 divided by Dextran. FIRE LUT is applied with a LUT calibration bar shown on the side. Quantification panels (**A’, B’**) show a plot of FluoZin-3/Dextran ratio of Ctrl (in black) and EGTA or *poky* (in magenta) as a function of time. At the sampled time resolution of 0.5 - 6 min, significant barrier breaches are detected in EGTA-treated or *poky* mutant embryos. Note that the intensity for the FluoZin-3 image of EGTA at 64’ and *poky* at 36’ was decreased for better display. Ctrl (EGTA): N = 4, n = 7; EGTA: N = 3; n = 6. Ctrl (*poky*): N = 3, n = 5; *poky*: N = 3; n = 5. N, number of independent experiments; n, number of embryos. Scale bars, 10 μm. See also **Movie 1**.

Finally, we examined whether ZnUMBA is applicable for cultured cells. For this purpose, we used MDCK II cells, which are one of the most commonly used epithelial cell lines for evaluating the structure and function of the TJ. Since wild-type (WT) MDCK II cells express cation-permeable claudin-2 (cldn2) (Amasheh et al., 2002; Furuse et al., 2001; Tokuda and Furuse, 2015), we used cldn-2-knockout (KO) MDCK II cells (Saito et al., 2021) (**Fig. S3, S4**). To observe cell sheets with a confocal microscope, the cells were cultured on the bottom-side surface of a Transwell filter (**Fig. 2C**). The bottom-side culture setup does not inhibit barrier formation compared with the normal top-side culture setup, although it takes 1-2 days longer to reach the full barrier function, and the TER value at plateau was slightly higher than the top-side culture conditions (**Fig. S5**). After the cells form a cell sheet, apical and basal media were replaced with ZnCl_2_- and FZ3-containing solutions (**Fig. 2C**). To test whether ZnUMBA can detect paracellular permeation of ions, WT MDCK II cells were mixed with cldn2-KO cells (**Fig. 5A**). To distinguish WT cells from cldn2-KO cells, nuclear localization signal-conjugated green fluorescent protein (GFPnls) was stably expressed in the WT cells (**Fig. S4**). Before the addition of FZ3, GFPnls alone was visualized in the green channel (**Fig. 5A**). Upon addition of FZ3, the FZ3 intensity at the cell-cell junctions between WT cells labeled with GFPnls drastically increased, whereas the signal between cldn2-KO cells remained modest (**Fig. 5A**), indicating that ZnUMBA can detect global changes in barrier function in MDCK II cells. Next, to evaluate whether ZnUMBA can be used to detect local breaches, we used angulin-1 (ang1)-KO cells. Angulin-1 localizes specifically at tTJs in MDCK II cells (Higashi and Chiba, 2020; Higashi and Miller, 2017), and loss of ang1 results in impaired barrier function at tTJs (Masuda et al., 2011; Sugawara et al., 2021). We established ang1-KO cells from the cldn2-KO clone and labeled them with GFPnls (**Fig. S3, S4**). When the ang1/cldn2-double KO (dKO) cells were mixed with cldn2-KO cells, FZ3 signal appeared preferentially at tricellular contacts of ang1/cldn2-dKO cells compared with those of cldn2-KO cells (**Fig. 5B**), indicating that ZnUMBA can detect local TJ barrier disfunction.

**Figure 5.**
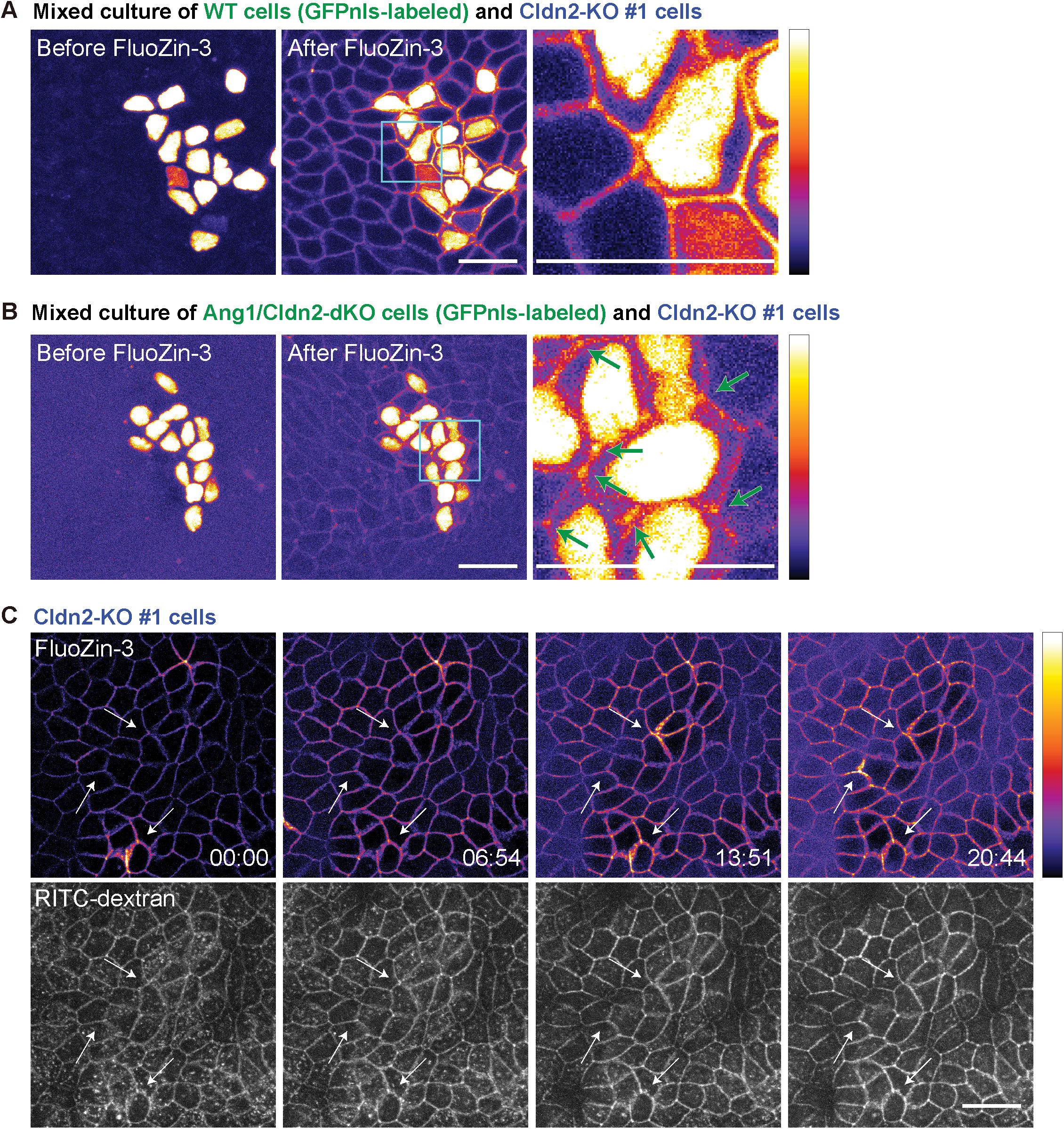
ZnUMBA detects transient local barrier defects in MDCK II cultured epithelial cells. **A.** ZnUMBA of WT cells and Cldn2-KO cells. GFPnls-labeled WT MDCK II cells and Cldn2-KO MDCK II cells were mixed and co-cultured on the bottom surface of the filter. Images before (left) and after (middle) FluoZin-3 addition are shown. FIRE LUT is applied. Right panel is the magnified view of the region outlined by a rectangle in the middle panel. Note that the cell-cell junctions between WT cells labeled with nuclear GFP have higher signal compared with those between Cldn2-KO cells. **B.** ZnUMBA of Ang1-KO cells in the Cldn2-KO background. GFPnls-labeled Ang1/Cldn2-dKO MDCK II cells and Cldn2-KO MDCK II cells were mixed and co-cultured. Note that tricellular junctions between Ang1/Cldn2-dKO cells have higher signal compared with the bicellular junctions (green arrows). **C.** Time-lapse images of ZnUMBA using Cldn2-KO cells. RITC-dextran (lower panels, white) is added to the ZnCl_2_ solution to visualize the basal compartment of paracellular space. FIRE LUT is applied to the FluoZin-3 channel. Note that ZnUMBA signal fluctuates over time whereas RITC-dextran signal remains unchanged. Bars, 20 μm. See also **Movie 2**.

We also tested performing ZnUMBA with live-imaging of MDCK II cells. In the cldn-2-KO cell sheet, ZnUMBA could also detect naturally-occurring leaks at cell-cell boundaries (**Fig. 5C, Movie 2**). In contrast with the spot-like FZ3 signals detected at naturally-occurring leaks in *Xenopus* embryos (Stephenson et al., 2019), the signals appeared to spread along the cell-cell boundaries across the vertices in MDCK II cell sheets. This difference might be because the Zn^2+^ ions can diffuse and spread in the intercellular spaces more easily in MDCK II cell sheets compared with intact *Xenopus* embryos. The locally increased FZ3 signals disappear within minutes, suggesting that there is a molecular mechanism to detect and repair the TJ breaks in MDCK II cells. Taken together, these data indicate that ZnUMBA is applicable for use in cultured epithelial cells and can detect leaks, which are either naturally-occurring or caused by loss of TJ components.

## Discussion

Here, we build upon recent publications utilizing the ZnUMBA method we developed (Chan et al., 2019; Stephenson et al., 2019; Varadarajan et al., 2021) to further explain ZnUMBA, demonstrate its capabilities, and provide examples of its use. In gastrula-stage *Xenopus* embryos, we show that ZnUMBA can be used, in combination with laser injury of the junction, to measure the dynamics of barrier restoration. Additionally, we demonstrate that by quantifying the FZ3/Dextran ratio, ZnUMBA can be used in zebrafish embryos to reveal barrier breaches with high spatiotemporal precision and different dynamic patterns: rapid barrier breaches were detected by global application of EGTA, and gradual leakage was monitored by genetically disrupted barrier function in *poky* mutants. Finally, in cultured MDCK II epithelial cells, we demonstrate that claudin-2 KO MCDK II cells (Saito et al., 2021) are suitable for detecting transient local barrier leaks with ZnUMBA, whereas the WT MDCK II cell line is quite leaky, causing significant background signal. Furthermore, we show that leaks at tTJs are detected in an angulin1/claudin-2 dKO MDCK II cell line. Together, these data underscore the utility and capabilities of ZnUMBA for detecting dynamic barrier breaches in a variety of systems.

Existing methods to measure barrier function, including TER, tracer permeability assays, and even sandwich assays, lack the ability to provide information about transient, local permeability changes caused by molecular perturbations, the addition of small molecules or peptides, or changes in cell shape or tissue mechanics. TER and tracer permeability assays provide measures of barrier function averaged across a population of cells, but no information about the location of barrier breaches, while sandwich assays provide subcellular information, but only a snapshot in time. In contrast, ZnUMBA allows analysis of subcellular epithelial permeability dynamics with high spatiotemporal resolution.

The existing methods to analyze barrier function generally involve analysis of cultured epithelial cells since they allow for easy perturbation and quantification of junctional properties; however, aspects of TJ structure and remodeling in cultured cells may not be fully representative of intact *in vivo* systems. Recently, variations on tracer permeability assays have been applied in order to monitor barrier function *in vivo*, aiming to measure local and/or dynamic TJ permeability in mouse models or even in human patients with intestinal barrier function diseases. For example, in mice orally gavaged with FITC-labeled dextran then later sacrificed, serum FITC fluorescence can be measured as a global readout; moreover, overlays of FITC fluorescent signal in a colon section and hematoxylin and eosin (H&E) staining of the same section can reveal information about local sites of barrier damage (Cario et al., 2007; Furuta et al., 2001). In a more precise but technically challenging *in vivo* murine method called exteriorized intestinal loop (iLoop), a well-vascularized exteriorized intestinal segment is made, the iLoop lumen is then injected with FITC-dextran, and intestinal permeability is assessed after an incubation period by quantifying fluorescence in blood serum (Boerner et al., 2021). iLoop offers an advantage over the oral gavage delivery of fluorescent tracer in that it allows researchers to study the permeability properties of specific localized areas in the intestine (terminal ileum or proximal colon) that are commonly involved in inflammatory bowel disease (IBD). Additionally, various reagents can be injected into the iLoop lumen, such as chemokines, cytokines, bacterial pathogens, toxins, antibodies and therapeutics to examine how these affect barrier function (Boerner et al., 2021). Perhaps in the future, iLoop could be combined with ZnUMBA to measure local and dynamic changes in TJ permeability. In human patients with IBD, confocal laser endomicroscopy (CLE) utilizes intravenous fluorescein injection and a confocal fluorescence microscope is incorporated in an endoscope in order to visualize the leakiness of intestinal epithelia. This technique has shown that there are particularly leaky areas of the intestinal epithelium *in vivo*, and these correlate with disease severity (Lim et al., 2014; Rasmussen et al., 2015).

In developing *Xenopus*, zebrafish, or other externally-developing model organism embryos, simple epithelia coat the surface of the embryo and are easily accessible for microscopy, making them well suited for live imaging of barrier function. In *Xenopus*, we have attempted visualizing the penetration of fluorescent molecules such as Alexa Fluor 488-Dextran (3,000 Da) (Reyes et al., 2014) or fluorescein (332 Da) (Higashi et al., 2016) across TJs by live imaging. Reyes et al. (2014) found that Anillin knockdown results in increased depth of penetration of Alexa Fluor 488-Dextran; however, this likely reflects cell shape change (apical doming) caused by disruption of junctional actomyosin rather than increased permeability because dextran did not detectably penetrate to the basal compartment (Reyes et al., 2014). Higashi et al. (2016) examined epithelial barrier function during cytokinesis by directly imaging fluorescein applied to the apical surface of *Xenopus* embryos. Under control conditions, it was not possible to detect the tracer beyond the TJ, even at the contractile ring, a site that promotes a major cell shape change, challenging junction integrity (Hatte et al., 2018; Higashi et al., 2016).

The variations on tracer permeability assays used for monitoring barrier function *in vivo* discussed above all suffer from a lack of specific signal increase at sites of local barrier leaks. Thus, a local, transient leak in the barrier would be difficult to detect against the background signal. Therefore, with ZnUMBA, we sought to develop a more sensitive barrier assay with minimal background, in which a breach of the TJ results in strongly increased fluorescence. Advantages of ZnUMBA with respect to existing techniques include: 1) FZ3 is a fluorogenic dye, and its fluorescence intensity increases ~50-fold specifically at barrier breach sites where FZ3 and zinc come in contact; 2) ZnUMBA is a live microscopy-based assay that can provide quantitative information about barrier function both at the tissue scale and subcellular level; 3) ZnUMBA is capable of revealing localized, transiently-occurring barrier leaks, thus identifying spatially where barrier breaches occur with respect to cell morphology or tissue architecture; and 4) ZnUMBA provides high temporal resolution, detecting changes in barrier function over time.

Despite its clear advantages, ZnUMBA also presents several limitations. First, embryo viability can be affected by high concentrations of ZnCl_2_; however, at lower concentrations of ZnCl_2_, the assay works effectively to detect barrier breaches, and does not cause phenotypic effects. Furthermore, exposing epithelia acutely to Zn^2+^ when performing ZnUMBA reduces the possibility that Zn^2+^ could affect barrier function, as different studies have reported that Zn^2+^ can either enhance or diminish epithelial barrier function (Shao et al., 2017; Wang et al., 2013; Xiao et al., 2018). As ZnUMBA is adapted for use in other systems, it will be important to empirically determine an appropriate concentration range of ZnCl_2_ and expose epithelial cells to the lowest dose of ZnCl_2_ for the shortest amount of time possible. Second, ZnUMBA can suffer from steadily increasing background FZ3 fluorescence over time. In *Xenopus* embryos, we found that decreasing the ZnCl_2_ concentration (1-2 mM), increasing the image acquisition exposure time, and adding Ca^2+^/EDTA along with FZ3 to sequester endogenous Zn^2+^ in the blastocoel improved background fluorescence and reduced noise. Nevertheless, the assay is still subject to gradually increasing FZ3 fluorescence over time; therefore, a method of normalizing signal to background is required for appropriate data interpretation. Finally, the selectivity of the pore pathway of paracellular permeability is dependent on which claudins are expressed (see below). Taken together, ZnUMBA offers many advantages for measuring dynamic changes in the TJ barrier at the subcellular level, and drawbacks to the assay can be overcome with careful experimental planning, controls, and normalization.

Some claudins including claudin-2, are pore-forming claudins and allow high volume of cations to cross the barrier, while restricting anions and other molecules. In the case of gastrula-stage *Xenopus* embryos, the most highly expressed claudins are claudin-6, claudin-7, and claudin-4 (Session et al., 2016), none of which are known cation pores (Gunzel and Yu, 2013). Interestingly, in the teleost fish group – especially in *Fugu rubripes*, but also in zebrafish – there has been extensive expansion of the claudin gene family, and several claudin genes have been identified that have no mammalian ortholog (Loh et al., 2004). This enrichment of claudin genes might have been due to an increased necessity for osmoregulation in fish species (Loh et al., 2004; Siddiqui et al., 2010). Some of the highly expressed genes in gastrula-stage zebrafish embryos include claudin-7b and claudin-8.2 (both anion-selective), claudin-b (orthologous to mammalian claudin-4, Na^+^-selective), and claudin-e (Siddiqui et al., 2010) and f (most closely related to claudins-3,4) (Kollmar et al., 2001). The broad expression pattern of several different zebrafish claudin genes with cation- or anion-selective functions supports the previously hypothesized osmoregulatory pressure in the case of zebrafish. There are two strains of MDCK cells that have similar TJ morphology but differ in the profile of claudins they express (Furuse et al., 2001). MDCK I cells do not express Claudin-2, whereas MDCK II cells do (Furuse et al., 2001; Tokuda and Furuse, 2015). MDCK II cells exhibit fewer clonal differences in junctional properties from one population to another, so they are generally preferred by TJ researchers (Matter and Balda, 2003). We found that WT MDCK II cells are quite leaky and not amenable for use with ZnUMBA. However, our data reveals that knocking down Claudin-2 in MDCK II cells makes them suitable for analysis by ZnUMBA.

Currently, ZnUMBA uses a commercially available dye, FZ3, applied to the basal side and ZnCl_2_ applied to the apical side of the barrier. These widely-available reagents and compatibility with conventional confocal microscopy make this assay accessible for potential use in diverse systems including developing model organisms, organ-specific setups like the exteriorized intestinal loops, 3D organoids, and 2D cell epithelial or endothelial cell culture. Additionally, because changes in TJ permeability underly multiple disease pathologies, and bacterial pathogens often target TJs, ZnUMBA could be used to better understand the mechanisms underlying these pathologies and to aid in drug discovery research by identifying compounds that modulate subcellular dynamics of barrier function. In the future, variations on the ZnUMBA method might be developed to expand the fluorescent dye color palette available or to measure size-specific permeability dynamics. Other fluorogenic dyes that increase significantly in fluorescence upon binding their target could be used instead of FZ3. For example, fluorogenic dyes that are ligands for self-labeling tags such as Halo tags (Zeng et al., 2019) would be a good option. In this case, the fluorogenic Halo dye would be added to the apical side of the epithelium, and a Halo-tagged basolateral transmembrane protein would be expressed in the cells, or recombinant Halo-tagged protein would be added to the basal side.

In closing, the development of ZnUMBA and its application in several systems including *Xenopus* (Stephenson et al., 2019; Varadarajan et al., 2021), mouse (Chan et al., 2019), zebrafish (this paper), and cultured epithelial cells (this paper) represents a powerful tool for detecting global and local dynamic changes in barrier function with spatiotemporal precision. We recommend that going forward, researchers should not only use traditional global, averaged measures of barrier function (TER, tracer permeability assays), but also measure local, dynamic changes in barrier function that can be detected with ZnUMBA. ZnUMBA has already revealed new information about subcellular dynamics of barrier function. New variations of ZnUMBA to meet different experimental needs should allow for broad application of this live imaging barrier assay to various epithelial contexts and will open the door to future discoveries.

## Materials and Methods

### ZnUMBA in *Xenopus* embryos *Xenopus embryo microinjections*

Laser injury experiments were performed on WT or albino *Xenopus laevis* embryos that had been microinjected with mRNA for Lifeact-mRFP (Bement et al., 2015) at the 1-, 2-, or 4-cell stage and were allowed to develop to gastrula stage at 15°C. Prior to imaging, 10 nl of 1 mM FluoZin-3 (FZ3, 847 Da, Thermo Fisher Scientific, Cat. F24194), 100 mM CaCl_2_, and 100 mM EDTA were microinjected into the blastocoel of stage 10-11 (Nieuwkoop and Faber) *Xenopus laevis* embryos. EDTA was used to reduce baseline levels of FluoZin-3 fluorescence from endogenous Zn^2+^, and equimolar Ca^2+^ was added to offset the potential effects of Ca^2+^ chelation by EDTA. Following microinjection, embryos were allowed to heal for a minimum of 5 minutes before being mounted in a slide containing 1 mM ZnCl_2_ in 0.1xMMR. Note: while we find it beneficial to reduce the time embryos are exposed to ZnCl_2_, we have found that embryos can be microinjected with FluoZin-3 and stored for several hours prior to imaging without negative consequences.

### Live microscopy and laser injury of junctions

Embryos were imaged using an Olympus FV1000 scanning confocal microscope with a 60x PlanApo objective (NA 1.4) and mFV10-ASW software. Six z-slices with a step size of 0.6 μm were collected every 3.5 seconds. A small circular ROI approximately the width of the junction (0.4-0.6 μm diameter) was placed roughly midway between two vertices, and a minimum of ten pre-injury frames were collected to establish a baseline. Injury was initiated manually by the user initiating the SIM scanner and 405 nm laser (100% power), and was stopped automatically after 5 seconds by the software. A maximum of three junction injuries was performed per embryo.

### Quantification of laser injury experiments

Quantification was performed in FIJI on summed z-projections with the assistance of custom macros and the LOI Interpolater tool in the Timelapse plugin. Every two to ten frames, a 1 μm wide segmented line was drawn over the junction from vertex to vertex, and the LOI Interpolater tool was used to draw lines in the remaining frames. Junction length, as well as mean intensity over the line of interest, were measured for FluoZin-3 and Lifeact-mRFP channels. FluoZin-3 and Lifeact-mRFP intensities were normalized by dividing the mean intensities of an injured junction by the mean intensities of a reference junction from the same movie. Junction length changes were normalized by subtracting the average junction length of the 10 frames prior to the injury from each junction length measurement.

### Zinc survival assay

Uninjected WT *Xenopus* embryos were allowed to develop to early gastrula stage at 15°C. Groups of 18-20 healthy embryos were incubated in 0.1xMMR plus the indicated concentration of ZnCl_2_ at room temperature (RT). Embryos were photographed prior to ZnCl_2_ exposure and after 1, 3, and 24 hours of ZnCl_2_ exposure. Abnormalities seen at gastrula stage (i.e., 1 and 3 hours) included small patches of lysing cells and exogastrulation. At the 24 hour time point, embryos were classified as abnormal if they were notably developmentally delayed or had developed abnormal growths. Embryos were classified as dead/lysing if they had cell material outside their body or if development had stopped and they were whitish in color.

### ZnUMBA in zebrafish embryos

#### Experimental model

Embryo collection from *Danio rerio* and staging of embryos were carried out as previously described (Kimmel et al., 1995; Westerfield, 2007). Embryos from AB strain were used as a control for *poky* mutant experiments, and AB or TL strains were used for EGTA experiments. For experiments, *poky* homozygous mutant fish were incrossed to gain maternal-zygotic *poky* mutants. Breeding of fish was performed at IST Austria zebrafish facility in line with local regulations and with the approval of the Ethic Committee of IST Austria regulating animal care and usage.

#### Zebrafish embryo injection

For mRNA transcription, SP6 mMessage mMachine Kit (Ambion) was used and mRNAs were injected via glass capillaries (30-0020, Harvard Apparatus, pulled by a needle puller P-97, Sutter Instruments) while mounted on a microinjection system (PV820, World Precision Instruments). Injections of 50 pg membrane-RFP (Iioka et al., 2004) with 0.2% Phenol red at 1-cell stage were performed as previously described (Westerfield, 2007). For FluoZin-3 injections, embryos were injected into the interstitial fluid with 100 μM FluoZin-3 together with 0.25 mg/ml Alexa Fluor 647 Dextran 10,000 MW (Invitrogen, #D22914) at 3 hpf (1k-cell stage).

#### Sample preparation for live imaging

Embryos were dechorionated. For live imaging of *poky* mutant experiments, embryos were mounted in 0.3% low melting point agarose (Invitrogen) on glass bottom dishes (MatTek). For live imaging of EGTA treatments, embryos were embedded into 0.3% low melting point agarose already containing 10-20 mM EGTA in calcium-free medium (isotonic Ringer’s Solution without Calcium: 116 mM NaCl, 2.9 mM KCl, 5 mM HEPES, pH 7.2 (Westerfield, 2007) and mounted in 4-chamber glass bottom dishes (MatTek) for imaging different conditions in parallel (EGTA addition vs. Danieau’s medium as control).

#### Live imaging of zebrafish embryos

High-resolution spinning disk confocal imaging was performed on a Zeiss Axio Observer Z1 microscope equipped with a 40x/1.2 W objective (C-APOCHROMAT, Korr UV-VIS-IR). Imaging was started at 4 hpf and within 30 min after addition of EGTA. The most superficial epidermal layer (enveloping layer, EVL) and deep cell interface were imaged by taking z-stacks between 10-25 μm at z-steps of 0.5-1 μm at acquisition times of 0.5-6 minutes.

#### Quantification of zebrafish ZnUMBA readout

FluoZin-3 fluorescence signal was normalized to the interstitial fluid abundance via Dextran 647 10,000 MW labeling. Maximum intensity projections (MIP) of the EVL and deep cell interface were generated to monitor the region in proximity to the barrier breach. Then, a region of interest with the size of 40 x 40 μm was selected, and FluoZin-3 mean intensity was divided by Dextran 647 mean intensity and plotted over time.

#### Titration of ZnCl_2_ concentration

In order to determine an appropriate concentration of ZnCl_2_, embryos were collected and exposed to different concentrations of ZnCl_2_ starting from around 3-4 hpf until 1-1.5 dpf in either dechorionated or chorionated conditions. ZnCl_2_ concentrations were 750 μM, 1 mM, or 2 mM. At higher concentrations (2 mM ZnCl_2_), a small fraction of embryos showed defects such lysis or abnormal development, while at lower concentrations (750 μM ZnCl_2_), embryos mimicked WT control viability rate.

### ZnUMBA in MDCK II cells

#### Antibodies

Rat anti-claudin-2 monoclonal antibody (mAb, clone 2D7) (Saito et al., 2021), rat anti-occludin mAb (clone MOC37) (Saitou et al., 1997) and rabbit anti-angulin-1 polyclonal antibody (pAb) were previously described. Rabbit anti-ZO-1 pAb (#61-7300), rabbit anti-claudin-2 pAb (#51-6100), rabbit anti-claudin-3 pAb (#34-1700), mouse anti-claudin-4 mAb (#32-9400), rabbit anti-tricellulin mAb (clone 54H19L38, #700191) and rabbit anti-ZO-2 pAb (#71-1400) were purchased from Thermo Fisher Scientific (MA, USA). Rat anti-ZO-1 (alpha+) mAb (clone R40.76; sc-33725), mouse anti-cingulin mAb (clone G-6; sc-365264) and goat anti-ZO-3 pAb (C-17; sc-11478) were from Santa Cruz Biotechnology (CA, USA). Rabbit anti-claudin-1 pAb (#18815) and rabbit anti-claudin-7 pAb (#18875) were form Immuno-Biological Laboratories (Gumma, Japan). Rabbit anti-occludin pAb (#LS-B2187) was from Lifespan (RI, USA). Rabbit anti-E-cadherin mAb (clone 24E10; #3195T) was from Cell Signaling Technology (MA, USA). Mouse anti-b-actin mAb (clone AC-15; #A1978) was obtained from Sigma-Aldrich (MO, USA). Rabbit anti-GFP pAb was from MBL (Nagoya, Japan).

For the secondary antibodies used in the immunofluorescence staining, Cy3-conjugated donkey anti-rat IgG pAb (#712-165-153), Alexa Fluor 488-conjugated donkey anti-rabbit IgG pAb (#711-545-152) and Alexa Fluor 647-conjugated donkey anti-rat IgG pAb (#712-605-153) were purchased from Jackson ImmunoResearch Laboratories (PA, USA). For immunoblotting, horseradish peroxidase (HRP)-linked sheep anti-mouse IgG pAb (#NA931V, GE Healthcare, CT, USA), HRP-linked goat anti-rabbit IgG pAb (#7074P, Cell Signaling Technology), HRP-linked goat anti-rat IgG pAb (#NA935V, GE Healthcare) and HRP-linked rabbit anti-goat IgG pAb (DAKO #P0449; Agilent, CA, USA) were used.

#### Cell culture

MDCK II cells were kindly provided by Prof. Mikio Furuse (National Institute for Physiological Sciences, Okazaki, Japan) and were maintained in DMEM (Sigma-Aldrich) supplemented with 10% fetal bovine serum (FBS; Sigma-Aldrich) at 37°C in a 5% CO_2_ incubator.

#### Immunofluorescence microscopy of cultured epithelial cells

MDCK II cells cultured on coverslips or Transwell filters with 0.4 μm pore size (#3401; Corning, NY, USA) were fixed with 1% formaldehyde at RT for 15 min. The cell membrane was permeabilized with 0.2% Triton X-100 in phosphate-buffered saline (PBS) at RT for 10 min. Then, the cells were blocked with 2% bovine serum albumin (BSA) in PBS at RT for 30 min, and were incubated with primary antibodies in PBS containing 0.2% BSA at RT for 1 hr, followed by secondary antibodies in PBS containing 0.2% BSA at RT for 30 min. The coverslips or filters were mounted with FLUORO-GEL II with DAPI (Electron Microscopy Sciences, PA, USA) and observed with an inverted laser scanning confocal microscope (FV1000; Olympus, Tokyo, Japan) with a 60x oil-immersion objective lens (UPlanSApo 60x; Olympus) at laser wavelengths of 405, 488, and 559 nm. Images were acquired with Fluoview ver. 4.2b software (Olympus) and processed with ImageJ 1.53a and Photoshop 2020.

#### Immunoblotting

MDCK II cells were lysed with sodium dodecyl sulfate (SDS) sample buffer (62.5 mM Tris-HCl, pH 6.8, 2% (w/v) SDS, 10% (v/v) glycerol, 5% (w/v) β-mercaptoethanol, 0.005% (w/v) bromophenol blue (BPB)), and proteins were separated in SDS-polyacrylamide gels (Fujifilm WAKO, Osaka, Japan). The proteins were transferred to PVDF membranes (Immobilon; Merck, Darmstadt, Germany) and blocked with 5% (w/v) non-fat dry milk in tris-buffered saline containing 0.1% (v/v) Tween-20 (TBST) at RT for 30 min. Then, the membrane was incubated with primary antibody diluted in TBST at 4°C overnight. The next day, the membrane was washed with TBST, incubated with secondary antibody at RT for 1 hr, and developed using the enhanced chemiluminescence method (ECL prime; GE Healthcare). The images were recorded using LAS4000 (GE Healthcare) and processed with Photoshop.

#### Establishment of Cldn2-KO MDCK II cells and Cldn2/Ang1-dKO MDCK II cells by CRISPR/Cas9

Cldn2-KO MDCK II cells were established as described previously (Saito et al., 2021). Specifically, 5 x 10^4^ MDCK II cells were transiently transfected with pSpCas9(BB)-2A-Puro (PX459) plasmids (Addgene, plasmid #62988, MA, USA) (Ran et al., 2013) encoding Cas9 and gRNAs for Cldn2 (sequences are shown in **Fig. S3**) using PEI-max (Polysciences, PA, USA). The next day, the cells were treated with 3 μg/mL puromycin (Sigma-Aldrich) for 1 day, and cells were sparsely reseeded onto a 10 cm dish. 7-10 days later, cell clones were picked, and aliquots of the cells were analyzed by genomic PCR using GoTaq DNA polymerase (Promega, WI, USA) and specific primers (F_C2, 5’-gatgccttcttgagcctgcttgtgg-3’; R_C2, 5’-agcaccttctgacatgatacagtgc-3’). The amplicons were cloned into pGEM-T-easy vector (Promega) using the TA-cloning method, and the DNA sequences were analyzed (Macrogen Japan, Kyoto, Japan). The cell clones where both alleles of the *claudin-2* gene locus are knocked out were chosen and further validated by immunofluorescence staining and immunoblotting. Likewise, Cldn2/Ang1-dKO cells were established using Cldn2-KO MDCK II cells as a parental cell line and were screened using specific primers for the *angulin-1/lsr* gene locus (F1_A1, 5’-tgctcctcgtccacgttatttcc-3’; F2_A1, 5’-cctctttctcagcaccttgtgcgc-3’; R1_A1, 5’-ggatcagcagatcccggcctc-3’).

#### Generation of nuclear GFP-expressing cell lines

5 x 10^4^ MDCK II cells or Cldn2/Ang1-dKO cells were transfected using PEI-max with a plasmid (pCAG-nGFP) encoding GFP conjugated with 3x nuclear localization signals (DPKKKRKVRS) (Saito et al., 2021). The cell clones were selected with 200 μg/mL of G418 (Sigma-Aldrich) for ~10 days and were screened by fluorescence microscopy.

#### ZnUMBA with cultured epithelial cells

To culture the cells on the bottom of the Transwell filters (#3401, Corning, NY, USA), the filters were placed upside-down in a clean container, and 1 x 10^5^ cells resuspended in 300 μL of DMEM were placed on the top of the Transwell to cover the entire surface of the filter. Then, the container was placed in a CO_2_ incubator and incubated at 37°C overnight. The next day, the filters were put into the wells of a 12-well plate filled with 2 mL of DMEM, and 500 μL of DMEM was added into the filters. The medium was changed every day. When the TER values reached a plateau, the filters were washed with Hanks’ balanced salt solution with Ca^2+^ and Mg^2+^ (HBSS, Gibco #14025-092; Thermo Fisher Scientific). A glass-bottom dish was placed on a oil-immersion 60x lens (UPlanSApo) of an inverted laser-scanning confocal microscope (FV1000) and 500 μL of HBSS containing 2 mM ZnCl_2_ was put on the center of the dish. Then, 200 μL of 10 μM FluoZin-3 solution in HBSS containing 1 μM CaCl_2_-EDTA was added into the filter, and the filter was placed on the glass bottom dish. Ca-EDTA chelates the excess Zn^2+^ in the basal compartment without sequestering Ca^2+^, which is required for the maintenance of cellcell junctions. The fluorescence signal of FluoZin-3 was detected using a 488-nm laser and the filter set for GFP. The images were recorded using Fluoview ver. 4.2b software and processed with ImageJ and Photoshop.

#### Transepithelial electric resistance (TER) measurement of cultured epithelial cells

MDCK II cells were cultured on the top or bottom surface of Transwell filters. The alternating-current (AC) electric resistance between the apical and basal compartments was measured each day using a volt-ohm meter Millicell ERS-2 (EMD Millipore, MA, USA). The electric resistance of a blank filter was subtracted from each measurement, and then the culture area of the Transwell filter (1 cm^2^) was multiplied to calculate the unit area resistance.

## Acknowledgements

This work was funded by NIH grant R01GM112794 to A.L.M and by Grants-in-Aid for Scientific Research 21K06156 from the Japanese Society for the Promotion of Science and the Grant Program for Biomedical Engineering Research from the Nakatani Foundation to T.H. and by funding from the European Union (European Research Council advanced grant 742573) to C.- P.H.

## Author Contributions

Conceptualization: T.H., R.E.S., C.S., and A.L.M; Formal analysis: T.H., R.E.S., C.S.; Funding acquisition: T.H., A.L.M, C-P.H., and H.C.; Investigation: T.H., R.E.S., C.S., K.H.; Methodology: T.H., R.E.S., C.S.; Project administration: A.L.M; Visualization: T.H., R.E.S., C.S.; Writing – original draft: T.H., R.E.S., C.S., and A.L.M; Writing – review & editing: all authors.

---

**Supplemental Figure 1.**
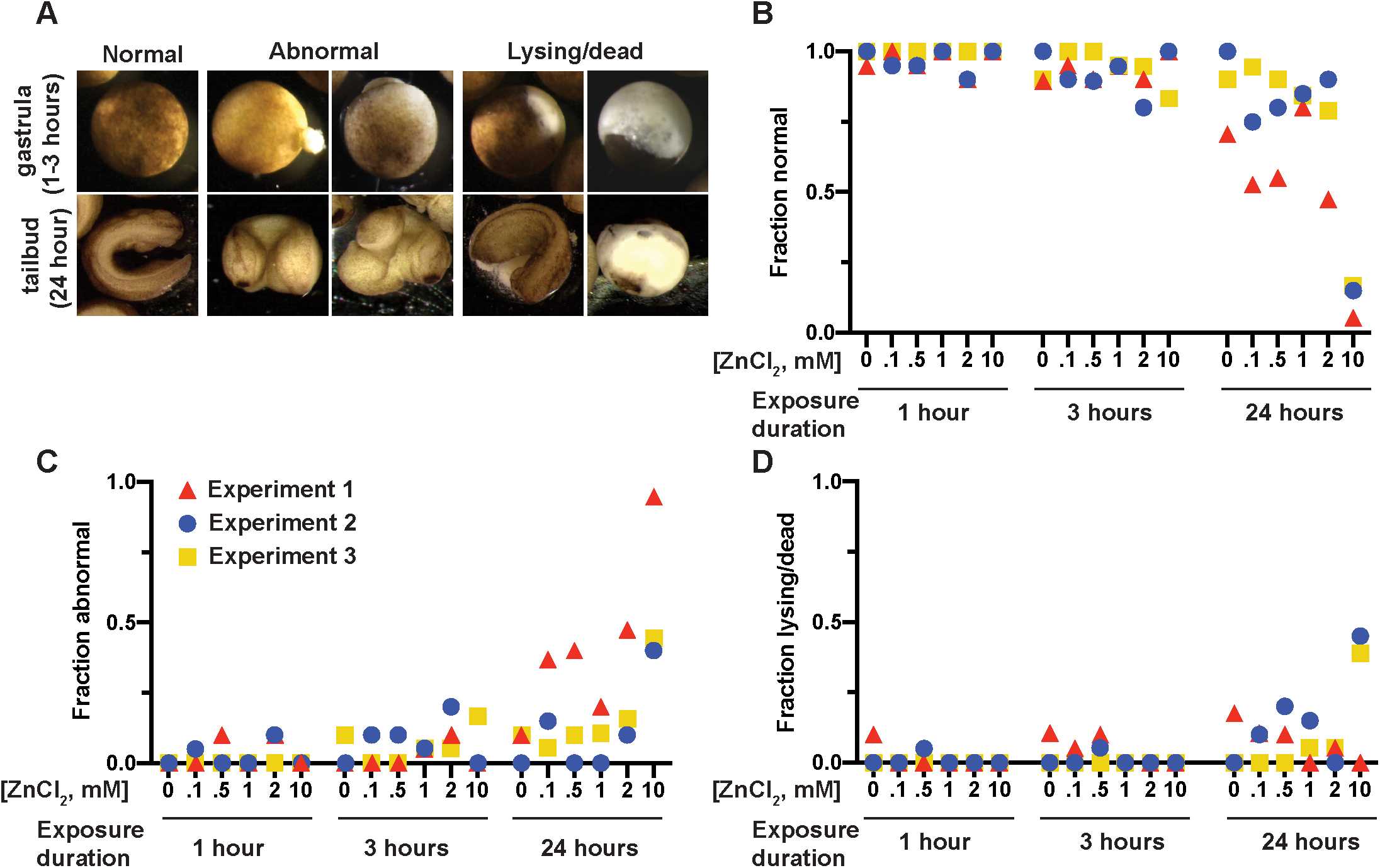
The effect of ZnCl_2_ on *Xenopus* embryo survival and development is time- and concentration-dependent. Gastrula-stage *Xenopus laevis* embryos were incubated at room temperature in 0.1xMMR containing the indicated concentration of ZnCl_2_ (from 0 mM indicated as control to 10 mM ZnCl_2_). At 1, 3, and 24 hours post-exposure, embryos were photographed and categorized as developing normally, developing abnormally, or lysing/dead. **A.** Examples of embryos classified as normal, abnormal, and lysing/dead (see Materials and Methods for more detail). **B-D.** The fraction of embryos classified as normal (**B**), abnormal (**C**), or lysing/dead (**D**) at the indicated ZnCl_2_ concentration and exposure duration. Points are color-matched by experiment. 3 independent experiments were performed with 18-20 embryos per ZnCl_2_ concentration.

**Supplemental Figure 2.**
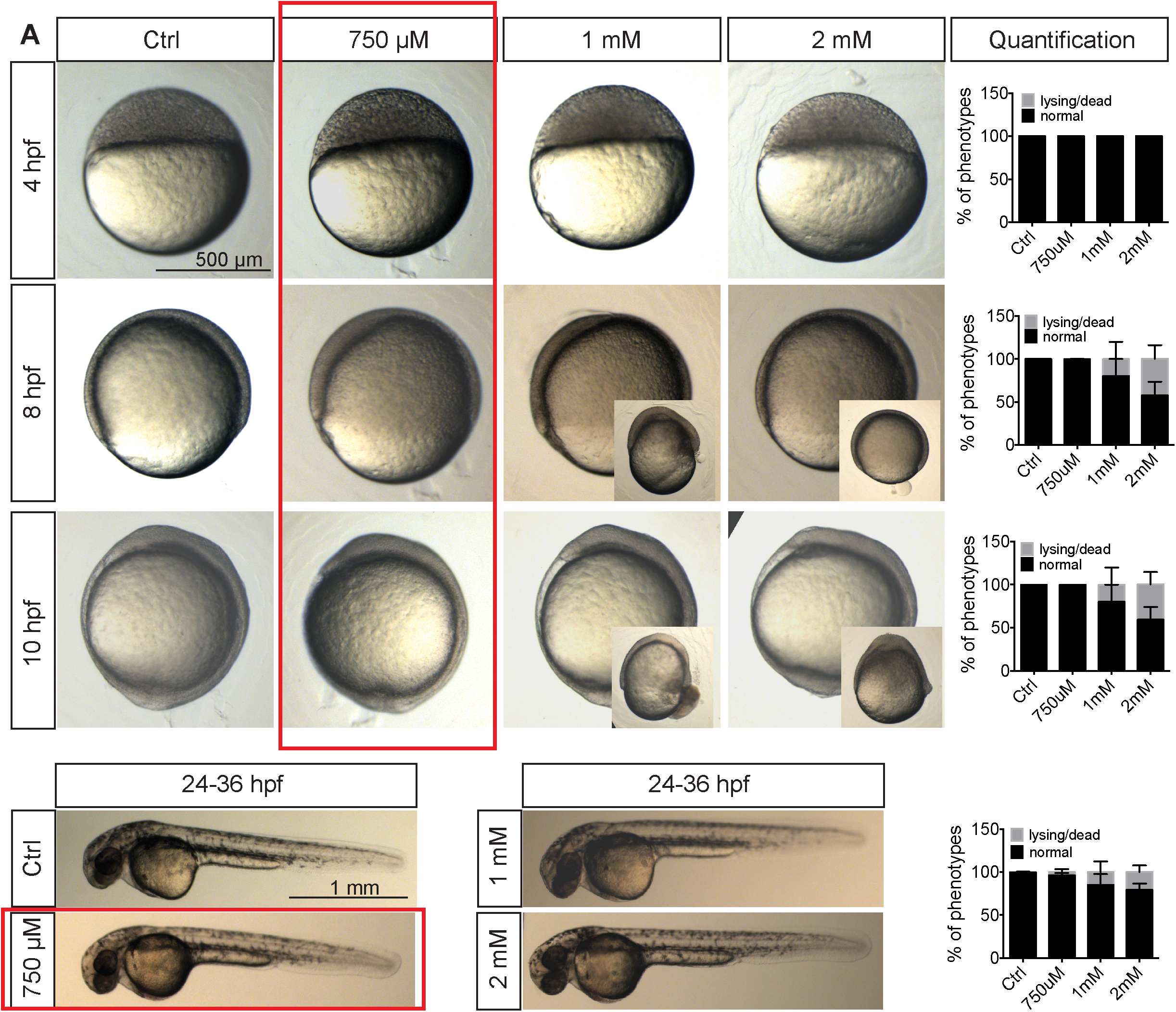
The effect of ZnCl_2_ on zebrafish embryo survival. **A.** Blastula-stage embryos were incubated at room temperature in Danieau’s medium containing the indicated concentration of ZnCl_2_ (from 0 mM indicated as control to 2 mM ZnCl_2_). At 4,8, 10, and 24-36 hpf, snapshots of embryos were taken and categorized as developing normally or lysing/dead. The insets in the snapshots indicate an example image of a lysing or dying embryo, while the large images show the main class of normal phenotypes. On the right column, the percentage of phenotypes categorized as normally developing or lysing/dead embryos is quantified. The selected condition (750 μM ZnCl_2_) is highlighted with a red rectangle. N > 3; n > 100. N, number of independent experiments; n, number of embryos. Scale bar, 500 μm (upper panel), 1 mm (lower panel).

**Supplemental Figure 3.**
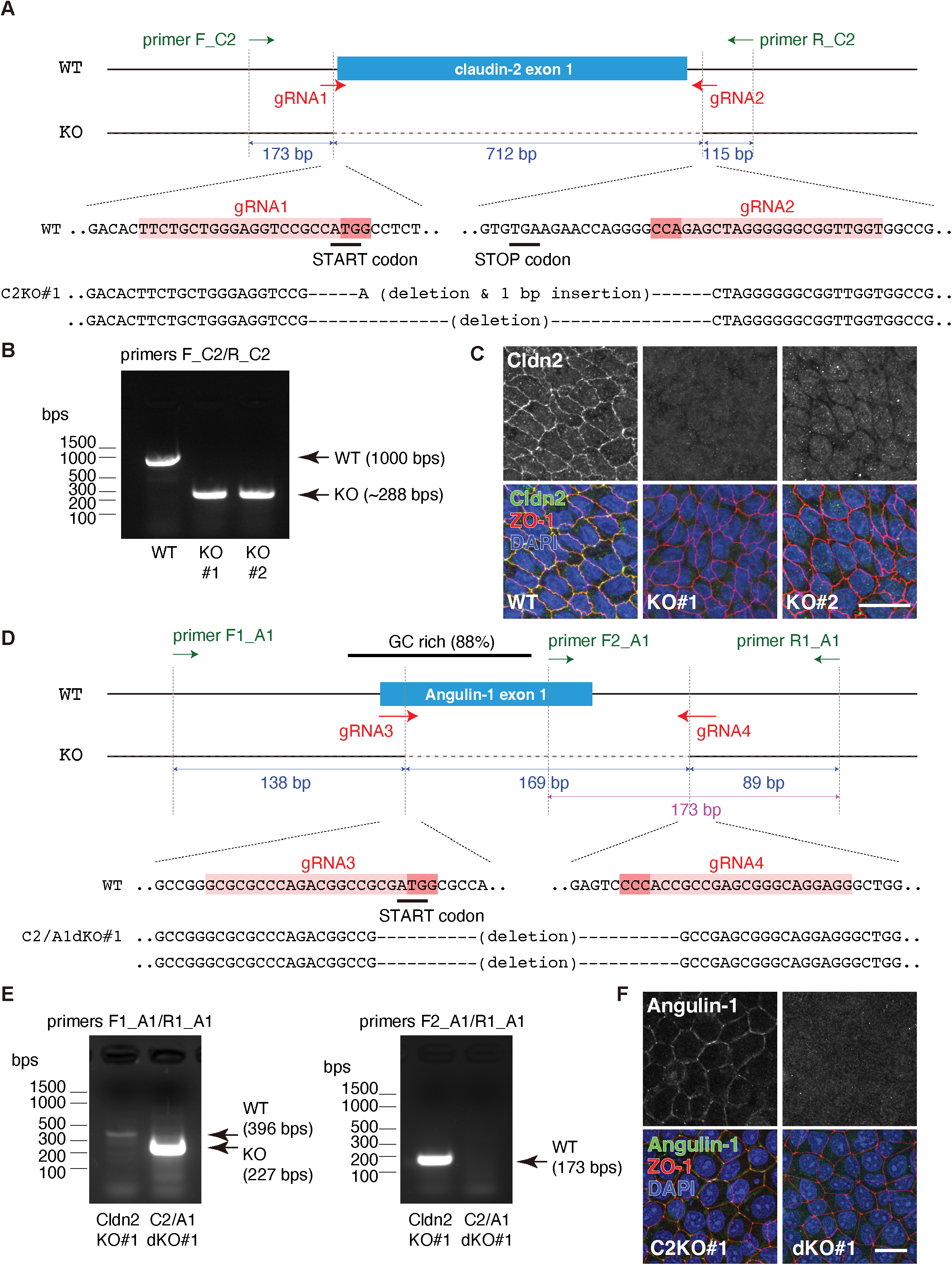
Generation of Cldn2-KO and Ang1/Cldn2-dKO MDCK II cells. **A.** Schematic of Cldn2 gene knockout. gRNA1 and gRNA2 are designed around the first ATG and in the 3’-untranslated region (UTR), respectively. Efficient knockout is confirmed by PCR using primer F_C2 and primer R_C2. Sequences of both alleles are shown. **B.** Genomic PCR result of C2KO#1 cells (newly established) and C2KO#2 cells (previously reported) (Saito et al., 2021). **C.** Immunofluorescence staining of WT, Cldn2-KO#1 and Cldn2-KO#2 cells. Cells were stained with anti-Cldn2 mAb (green), anti-ZO-1 pAb (red), and DAPI (blue). Bar, 20 μm. **D.** Schematic of Ang1 gene knockout. gRNA3 and gRNA4 were used to excise the exon 1 which contains the first ATG. The positions of primers F1_A1, F2_A1 and R1_A1 used for screening are shown. **E.** Genomic PCR results of Cldn2-KO#1 cells and Cldn2/Ang1-dKO#1 cells. Since most of the exon 1 of Ang1 gene is highly G/C-rich, amplification of fragments using primer F1_A1 and primer R1_A1 was very weak when the allele is intact (left, Cldn2-KO#1 lane). PCR using primer F1_A2 and primer R1_A1, which amplifies the fragment outside of the G/C rich region, further validated the efficient knockout of Ang1 gene in the Cldn2/Ang1-dKO cells. **F.** Immunofluorescence staining of Cldn2-KO#1 cells and Cldn2/Ang1-dKO cells. Cells were stained with anti-Angulin-1 (green), anti-ZO-1 mAb (red), and DAPI. Bar, 20 μm.

**Supplemental Figure 4.**
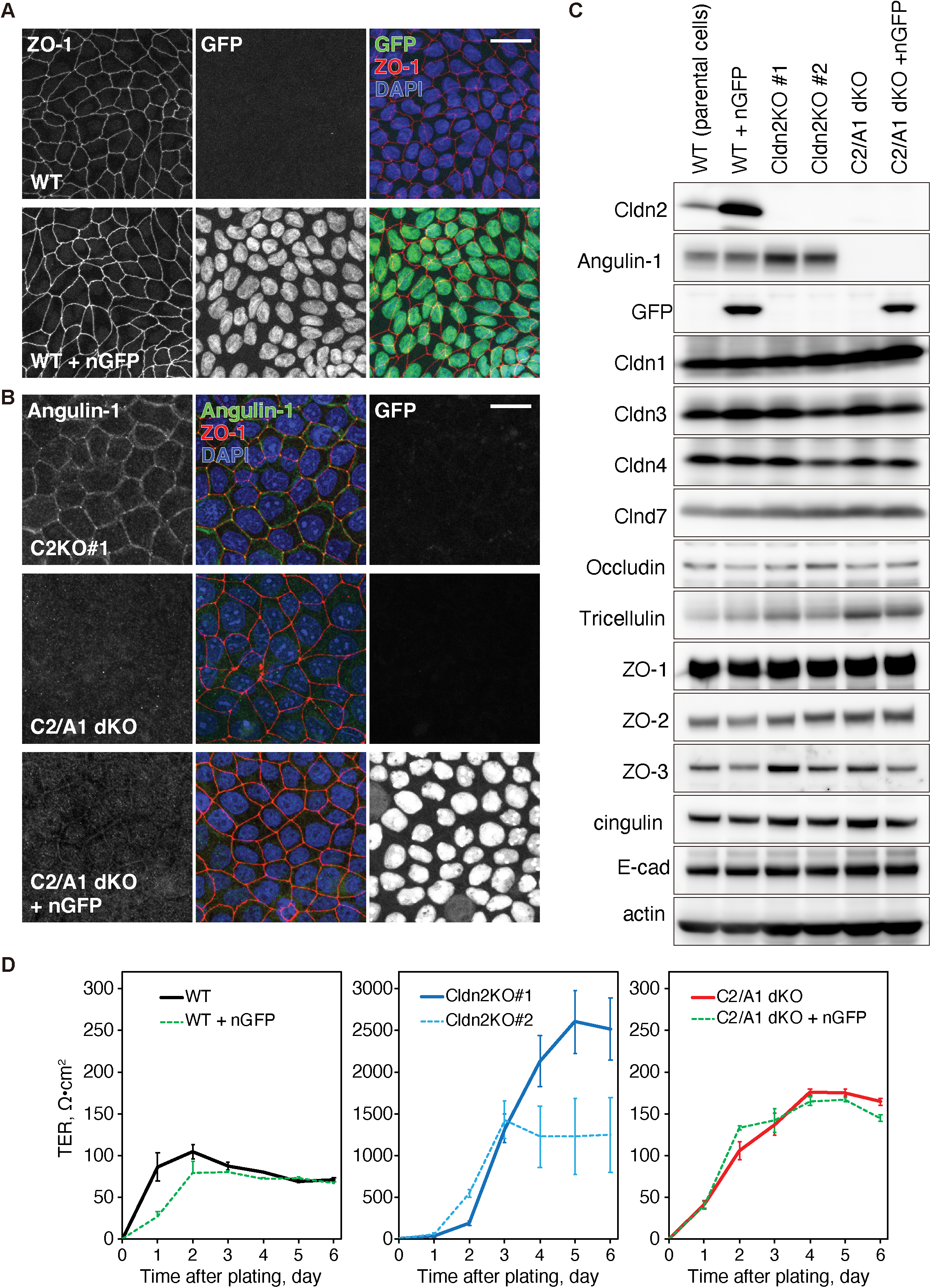
Characterization of TJs and global barrier function in Cldn2-KO and Ang1/Cldn2-dKO MDCK II cells. **A-B.** Immunofluorescence staining of WT and WT+nGFP cells (**A**) and Cldn2-KO#1, Cldn2/Ang1-dKO, and Cldn2/Ang1-dKO+nGFP cells (**B**). Cells were stained for ZO-1 (red) and DAPI (**A**) or Angulin-1 (green), ZO-1 (red), and DAPI (blue). Nuclear GFP signal is shown in green (**A**) or white (**B**). Bar, 20 μm. **C.** Immunoblotting of MDCK II cells used in this study. **D.** TER measurement of MDCK II cell clones. n = 9.

**Supplemental Figure 5.**
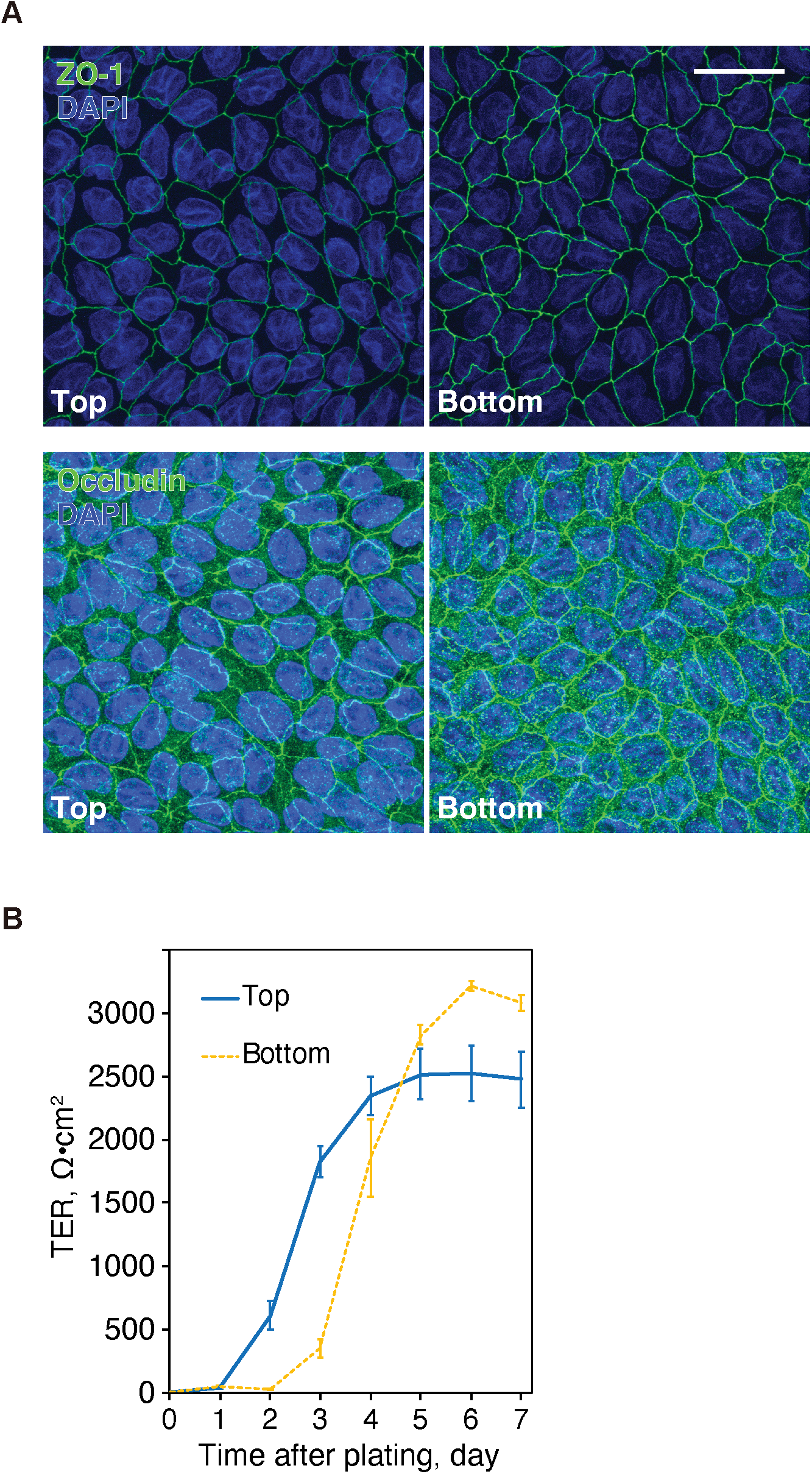
Characterization of Cldn2-KO MDCK II cells cultured on the bottom surface of a transwell filter. **A.** Immunofluorescence staining of Cldn2-KO MDCK II cells cultured on the top (left panels) or bottom (right panels) surface of a transwell filter. Cells were stained for ZO-1 (green, upper panels) or occludin (green, lower panels) and DAPI (blue). **B.** TER measurement of Cldn2-KO MDCK II cells cultured on the top or bottom surface of the filter. n = 10. Bar, 20 μm.

**Movie 1. ZnUMBA in EGTA treated zebrafish embryos.**

Time-lapse fluorescence movie of embryos labeled with membrane RFP in red to detect EVL and deep cells, FluoZin-3 in green, and Dextran-647, 10,000 MW in magenta. Representative control embryos incubated in Danieau’s medium (upper panel) and embryos incubated in Calcium free isotonic Ringer’s solution with 10-20 mM EGTA (lower panel). Movie was started at 4 hpf. Scale bar, 10 μm. Time interval, 2 min.

**Movie 2. ZnUMBA of Cldn2-KO MDCK II cells.**

ZnUMBA is conducted using Cldn2-KO cells. FIRE Lookup Table (LUT) is applied to the FluoZin-3 channel. RITC-dextran was added to the basal compartment to visualize the basolateral paracellular spaces. Time is indicated as min:sec. Note that transient localized increase and decreases of FluoZin-3 signal are observed.

